# A scalable phenotyping approach for female floral organ development and senescence in the absence of pollination in wheat

**DOI:** 10.1101/2022.04.01.486528

**Authors:** Marina Millan-Blanquez, Matthew Hartley, Nicholas Bird, Yann Manes, Cristobal Uauy, Scott Boden

## Abstract

In the absence of pollination, female reproductive organs senesce leading to an irrevocable loss in the reproductive potential of the flower and directly affecting seed set. In self-pollinating crops like wheat (*Triticum aestivum*), the post-anthesis viability of the unpollinated carpel has been overlooked, despite its importance for hybrid seed production systems. To advance our knowledge of carpel development in the absence of pollination, we created a relatively high-throughput phenotyping approach to quantify stigma and ovary morphology. We demonstrate the suitability of the approach, which is based on light microscopy imaging and machine learning, for the detailed study of floral organ traits in field grown plants using both fresh and fixed samples. We show that the unpollinated carpel undergoes a well-defined initial growth phase, followed by a peak phase (in which stigma area reaches its maximum and the radial expansion of the ovary slows), and a final deterioration phase. These developmental dynamics were largely consistent across years and could be used to classify male sterile cultivars, however the absolute duration of each phase varied across years. This phenotyping approach provides a new tool for examining carpel morphology and development which we hope will help advance research into this field and increase our mechanistic understanding of female fertility in wheat.

## Introduction

The fertilisation of the pistil by a pollen grain is a vital event in the life cycle of a flowering plant as it contributes to the reproductive fitness of a species. In grasses, the pistil (or carpel) typically consists of an ovary bearing two styles densely covered by a feathery and dry-type stigma [1, 2]. The stigmatic tissue plays a key role in successful fertilisation as it facilitates the interception and hydration of the pollen grain and mediates pollen tube growth into the stylodia branches towards the ovary containing the ovule [3, 4]. After successful fertilisation, the ovary undergoes growth and differentiation to develop into a grain. Under favourable growing conditions, the duration of carpel receptivity (or functionality) does not present a serious limitation to seed formation in self-pollinating species, such as wheat (*Triticum aestivum*) or rice (*Oryza sativa*). However, environmental stresses such as heat and drought [5-7] or the absence of viable pollen (e.g., male sterile cultivars used in hybrid breeding [8]) can affect normal seed set.

Female floral organs have developed a series of survival mechanisms to secure seed set in the absence of self-pollination by increasing the likelihood of receiving pollen from neighbouring male fertile plants. Indeed, this process (directly or indirectly) has been harnessed by breeders to produce hybrid seeds in crops like maize (*Zea mays*), rice, barley (*Hordeum vulgare*), and wheat. In maize, one of the survival strategies described to increase pollen capture is silk (i.e., stigma) emergence and elongation from the husk [9], while in wheat, the radial expansion of the unfertilised ovary pushes the floret open facilitating the access to airborne pollen, a phenomenon known as the “second opening” [10, 11]. However, if pollination still does not occur after a specific time, which varies between species [12, 13], a series of developmental processes leads to the senescence of the floral organs and the irreversible loss of reproductive potential [14]. For example, in several plants, the loss of papilla integrity has been regarded as one of the primary symptoms indicating the end of the floral receptive period and stigma senescence which is often manifested by the shrunken appearance of the stigma [11, 15, 16]. Similarly, the unfertilised ovary undergoes a series of morphological changes that converges in the lignification of the epidermal cells and eventual collapse of the ovary walls [11, 14]. In many of these studies, the phenotypic characterisation of these processes is time-consuming and labour intensive and is, therefore, usually performed only under controlled growing conditions and on a small number of plants. These phenotyping approaches, although extremely informative, are often not conducive for translation into breeding targets where the screening of large germplasm sets is required.

In recent years, high-throughput phenotyping technologies have provided new opportunities to phenotype a diverse range of plant species at various scales, ranging from cellular to tissue and organ levels [17, 18]. For instance, machine learning based algorithms, like neural networks, have become an essential tool for reliably extracting morphological information and providing visual quantitative parameters of microscopy images. These approaches can be used in large-scale experiments, like those of crop breeding programmes, and provide a way to quantify the morphological changes of the developing carpel in the absence of pollination.

To advance our knowledge of carpel development in the absence of pollination, we developed a phenotyping approach for the quantification of stigma area and ovary diameter of field grown plants by combining light microscopy and machine learning. We focused on bread wheat carpels due to the current need to improve outcrossing rates in hybrid breeding programmes [19] and the lack of knowledge on the dynamics of stigma and ovary development among male sterile (MS) wheat cultivars under production conditions in the field. We applied our phenotyping approach to three MS cultivars during two consecutive field seasons to gain insights into genetic and environmental variation for these two florals traits and show that it is scalable to produce practical advances in breeding programmes. Finally, we outline lessons, challenges and the opportunities that this phenotyping approach offers to gain a more comprehensive understanding of female fertility in cereal crops such as wheat.

## Materials and Methods

### Germplasm

We used both spring and winter male sterile hexaploid wheat (*Triticum aestivum*) cultivars derived from commercial inbred lines. Cytoplasmic and nuclear male sterility (CMS and NMS) systems were used for the generation of the male sterile cultivars. BSS1 and GSS2 correspond to winter CMS cultivars while the winter cultivars 24511, 24512, 24516 and 24522, and the spring cultivars Jetstream, Alderon, BLA1, Mairra, Cadenza, Chamsin and BLA2 are NMS cultivars. We used MS cultivar 24516 as an example to illustrate the developmental dynamics of the unpollinated wheat stigma and ovary in Fig. 1. All CMS and NMS cultivars were provided by KWS Ltd (Thriplow, UK) and Syngenta (Whittlesford, UK), respectively.

**Figure 1.**
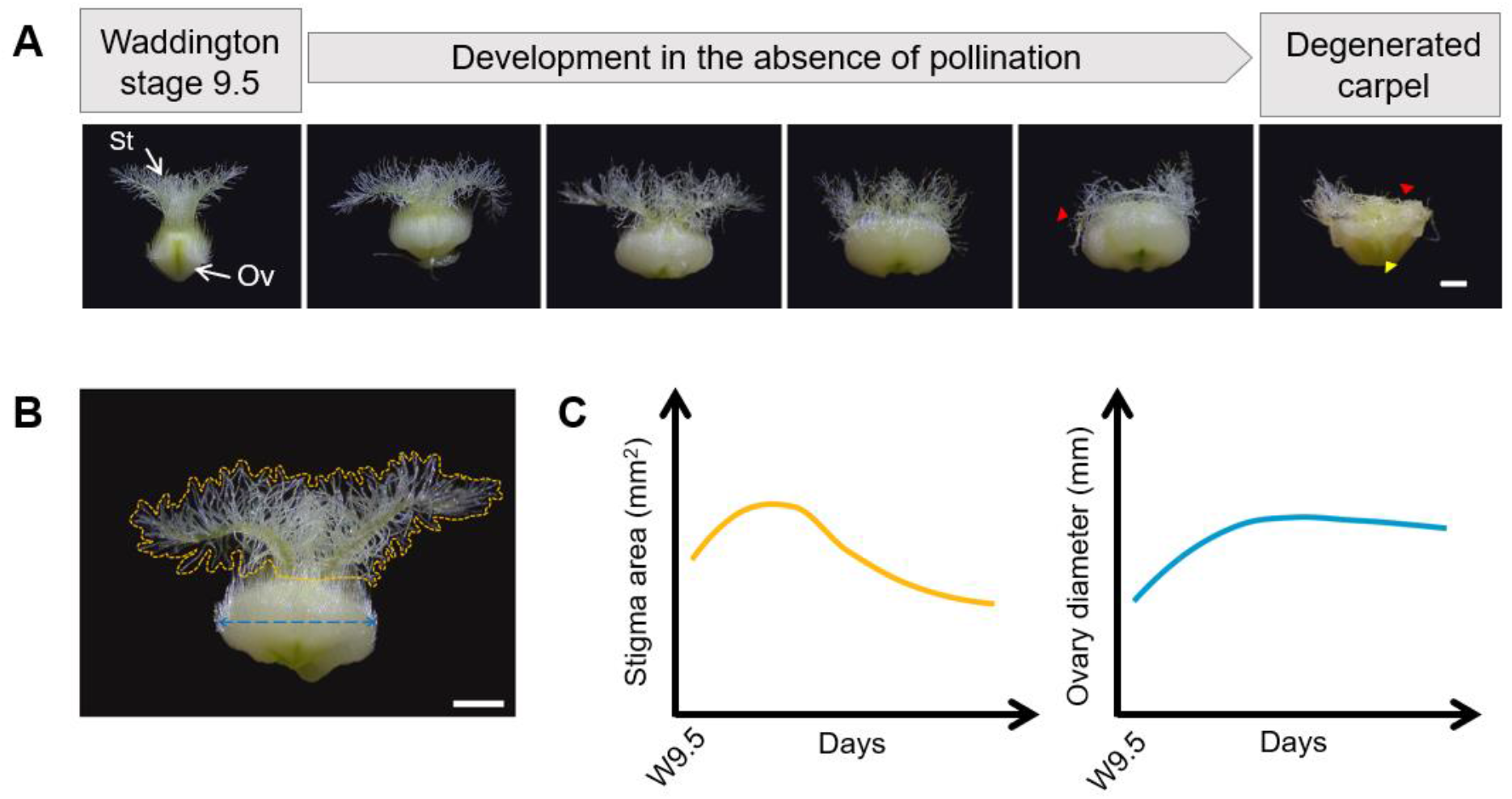
Representative stages of late carpel development in the absence of pollination. **(A)** Morphological changes observed in the stigma (St) and ovary (Ov) from Waddington stage 9.5 (approx. ear emergence) until complete degeneration of the carpel (from left to right). Arrowheads indicate regions of the stigma (red) and ovary (yellow) where symptoms of cell degeneration are visible. **(B)** Diagram illustrating the morphological traits of interest: the yellow line delimits the area covered by the stigmatic hairs, the blue line depicts the diameter of the ovary. **(C)** Representative growth trends observed for the stigma hair area (yellow) and ovary diameter (blue) in the absence of pollination for field grown plants. For the regression curves we have used a single exemplar male sterile cultivar (*n* = 10-30 carpels from 6 plants sampled at 8 timepoints). In B and C, scale bar = 1 mm.

To train the convolutional neural network (CNN) for the quantification of carpel traits we used a random sample of plants extracted from a set of the seven spring NMS cultivars. The selection criteria for the generation of the training set, however, were based on diversity of carpel images rather than on a per cultivar-based criteria.

We used four winter male sterile cultivars (24511, 24516, BSS1, GSS2) to characterise the effects of the fixative on stigma area and ovary diameter during the 2020 field season. Finally, for the multi-year field experiment performed to investigate the developmental patterns of the unpollinated carpel, we grew three winter male sterile cultivars (BSS1, 24512 and 24522) during two consecutive years.

### Glasshouse and field experiments

For the development of the stigma and ovary CNNs we grew between 10 to 20 plants per MS cultivar (Jetstream, Alderon, BLA1, Mairra, Cadenza, Chamsin and BLA2) in the glasshouse in 1 L pots under long day conditions (16 h light: 8 h dark) and hand dissected carpels from either the first, second or third tiller at various timepoints representative of the different carpel morphologies. We stored a random selection of the dissected carpels in a fixative solution of 95% ethanol and absolute acetic acid (75% v/v) and kept them at 4 °C until image acquisition (approx. a month after fixation). The cultivars selected are representative of carpel morphology diversity in wheat germplasm.

For the carpel development time course experiments, we grew plants at John Innes Centre Church Farm (Bawburgh, UK; 52°37’50.7” N 1°10’39.7” E) in a randomised complete block design with two replicates (plots) per MS cultivar in 2020, and three replicates in 2021 (Supplementary Figure 1).

To avoid unwanted cross-pollination, sterile rye barriers were grown surrounding the male sterile plots. To record environmental data, we placed data loggers (EasyLog USB, Lascar Electronics) next to the experimental plots at 50 cm height. Average temperature and relative humidity were measured every hour during the duration of the experiment (Supplementary Figure 2).

In both field seasons, we tagged main spikes when carpels in the outer florets (floret 1 and 2) of central spikelets reached Waddington stage 9.5 (W9.5 [20]). This is shortly after full ear emergence (Zadok’s growth stage 59 [21]). At the time of sampling, we cut individual tillers between the uppermost and penultimate internode and transported them in water to the laboratory for carpel dissection. We harvested four to seven carpels from the outer florets of central spikelets from two spikes per plot and timepoint. These timepoints were W9.5 and 3, 7, 13 and 18 days after W9.5 (DAW9.5), with the only exception that the 2021-time course was extended for 9 more days (i.e., 27 DAW9.5) in cases where the carpel was not completely senesced at 18 DAW9.5. Due to the limited availability of spikes tagged at W9.5, sample size for the extended timepoints varied from 2 to 6 spikes per timepoint. Carpels that needed fixation were placed in 2 mL Eppendorf tubes containing fresh fixative solution, as described above.

### Image acquisition and manual annotations

To generate the training set we used two different stereo microscopes equipped with an integrated camera for image capture (ZEISS Stemi 305 with a 1.2 Megapixel integrated colour camera; Leica MZ16 coupled with a Leica CLS100x white light source and a Leica DFC420 5 Megapixel colour camera). For the downstream experiments using the adapted CNNs we only used the ZEISS Stemi 305 since it is easier to operate and to transport to the field. Depending on the size of the carpel, we used different magnifications (from 1x to 4x) to ensure that the complete carpel was captured in the image (Supplementary Figure 3). We adapted illumination and time of exposure to each image to ensure high contrast between the carpel and the black background. To maintain the feathery structure of the stigma in fixed samples, we imaged the carpels submerged in a 70% ethanol solution using a Petri dish (one carpel per image). Images were stored as RGB jpeg files.

To evaluate the effect of the fixative on carpel morphology traits, we first imaged the carpels as non-fixed samples and we then placed them in the fixative solution (as described previously) for image acquisition at a later time. For manual annotations of stigma and ovary perimeters, we analysed the resulting images using open-source image processing package Fiji.

### Development of stigma and ovary CNNs

To carry out automated image annotation and measurement, we trained a CNN using the UNet design [22]. The network was implemented in the PyTorch framework [23], using the dtoolAI library [24]. The networks were trained on carpel jpeg images with manual annotations of the stigma perimeter (*n* = 86 images) and ovary perimeters (*n* = 121 images). These 207 images were randomly selected from a total of 1601 dissected carpels. The Dice coefficient [25] was used as a loss function, and weight updates were applied using the Adam optimizer[26].

The trained stigma network predicted a mask corresponding to the stigma location for each RGB input image. The stigma mask was used to calculate the stigma area directly by extracting the number of pixels. The trained ovary network predicted two masks for each RGB input image, one corresponding to the ovary location and the other to the stigma location. To determine the ovary diameter, the following algorithm was applied:

1. Determine the centroid of the predicted stigma mask.
2. Determine the centroid of the predicted ovary mask.
3. Take the perpendicular to the line drawn through the centroids.
4. Determine the two points where this line crosses the border of the predicted ovary mask.
5. Measure the length of this line, converting to physical units from the original input image scale.

We converted pixels (CNN output) to stigma area (mm^2^), and ovary diameter (mm) according to the scale bar used in each image. Implementation scripts and data are available at https://github.com/marina-millan/ML-carpel_traits.

### Statistical analyses and data visualisation

#### Evaluation of CNN performance

To evaluate differences in stigma area and ovary diameter between the manual and CNN annotations, we selected a random set of 60 microscopy images that were not used to train the networks (30 images of fixed carpels and 30 of non-fixed carpels). We divided the images into three different developmental stages based on the appearance of the carpel to account for all the possible carpel morphologies the algorithm could be exposed to. Stage 1 represents a young carpel (stigma and ovary still developing), stage 2 represents a fully developed carpel (widely spread stigma and enlarged ovary), and stage 3 includes visibly deteriorated carpels. We performed one-way analysis of variance (ANOVA) for each trait and sampling method, including “floral age” as the single factor (Supplementary Table 1). To measure the spatial overlap between the manual and CNN annotation, we calculated Dice similarity coefficients on the same set of images.

#### Carpel development time courses

For the quantification of stigma area and ovary diameter of fixed and non-fixed samples, a total of 666 and 634 images were annotated by the stigma CNN and ovary CNN, respectively, and used for subsequent analyses. We conducted three-way ANOVAs (fixative, timepoint, cultivar) to test the effect of the fixative on stigma area and ovary diameter and their interaction with floral age (timepoint) and cultivar (Supplementary Table 2). We include in the model block and spike information as random effects to account for the nested nature of the experimental design.

To generate the patterns describing stigma and ovary development for MS cultivars 24512, 24522 and BSS1, 294 and 520 images were annotated by the stigma and ovary CNNs, in 2020 and 2021, respectively, and used for downstream analyses. Next, we filtered out outliers following the interquartile range criterion and used loess smooth lines with a span value of 0.9 (i.e., width of the moving window when smoothing the data) and a 95% confidence interval. To have an estimate of the amount of growth the plants achieved during the different field seasons we calculated cumulative degree days using 0 °C as base temperature (according to Perry Miller et al. [27]) and average daily temperatures:

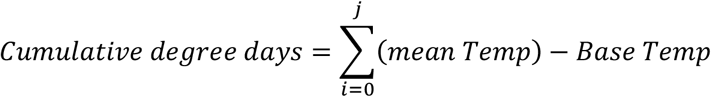

*i*: beginning of the temporal window considered (e.g., W9.5 date)

*j*: end of the temporal window considered (e.g., sampling date at 7 days after W9.5)

Statistical analyses and codes to model carpel and ovary development were conducted in RStudio Version 4.0.3 and are available at: https://github.com/marina-millan/ML-carpel_traits.

## Results

### Defining quantifiable parameters of late carpel development and senescence

To investigate the development of wheat carpels in the absence of pollination and determine parameters that correlate with key phases of its life cycle, we used nuclear and cytoplasmic male sterile plants. We imaged the unpollinated carpels of field-grown plants starting from Waddington stage 9.5 (W9.5, normally shortly after ear emergence; Fig. 1A) which corresponds with the most advanced developmental stage for an unpollinated carpel; W9.5 is only shortly before when male-fertile plants would reach anthesis (at W10) and release viable pollen on the receptive stigma surface. At W9.5, stigma branches are well elongated and spread outwards to generate the plumose architecture, while the unfertilised ovary shows a round shape (Fig. 1A). In subsequent timepoints after W9.5, the unfertilised ovary will gradually expand horizontally leading to the “second opening” of the floret (as previously described by Okada et al. and Molnár-Láng et al. [10, 11]). During this period of ovary growth, stigma branches continue growing and quickly curve away from each other, increasing the stigma surface area and contributing to the extrusion of unpollinated stigma outside the floret and thus, to the capture of airborne pollen. Towards the end of the time course, papilla cells of the stigma hairs start to lose turgor and become atrophied as the stigma degenerates (Fig. 1A, red arrowheads). Finally, the onset of stigma degeneration is followed by a slight and gradual decline in ovary radial size, causing the floret to close again [10].

To quantify the observed morphological changes in these parameters, we imaged unpollinated carpels and manually delineated the area covered by the stigma hairs and the diameter of the ovary using Fiji (Fig. 1B). We distinguished a developmental pattern for stigma area characterised by an initial bell curve shape followed by a gradual reduction in area indicative of tissue deterioration. The ovary diameter gradually increased and reached a plateau with little changes in the diameter thereafter (Fig. 1C). More importantly, these patterns appear to be quite consistent across different cultivars and replicates (see Results below). Together, these findings suggest that stigma area and ovary diameter are promising parameters to quantitively track the life cycle of the unpollinated carpel.

### Overview of the approach

The rapid and accurate phenotyping of large numbers of field-grown plants represents a challenge for plant researchers. Here, to accelerate research into female floral traits, we propose a phenotyping approach that can be implemented in the screening of large populations, such as those of pre-breeding programmes (Fig. 2). This approach provides a visualization and quantification toolbox for morphometric information of stigma area and ovary diameter. A summary of each step is provided below, and detailed descriptions can be found in the Materials and Methods section.

**Figure 2.**
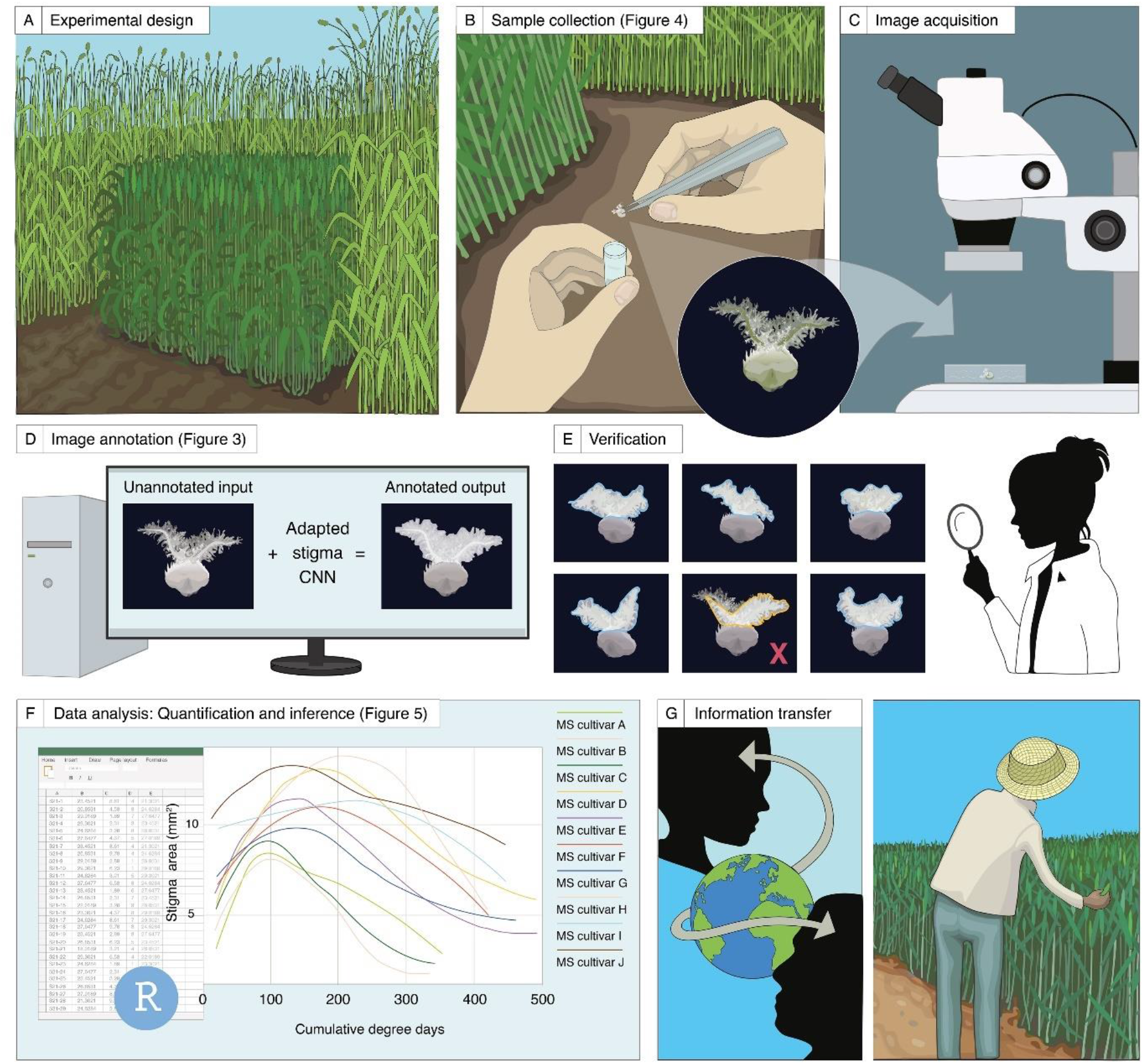
Schematic representation of the proposed phenotyping approach for the study of carpel development in the absence of pollination under field conditions. **(A-C)** Experimental design, sample collection and image acquisition. **(D-E)** Annotation of microscopy images and quantification of the stigma area (as an example, **D**) and verification of the CNN outputs **(E). (F)** and **(G)** illustrates the applicability of the phenotyping approach to enhance our understanding of the post-anthesis behaviour of unpollinated carpels.

### Experimental design, sample collection and image acquisition

As our main aim was to study carpels in the absence of pollination, the first step is to prevent cross-pollination of MS plants in the field. To achieve this, different strategies can be used. For example, in this study we grew sterile rye surrounding the experimental plots to create an effective pollen barrier from surrounding fertile plants (Fig. 2A and Supplementary Figure 1). When anthers from the outer florets of the central spikelets are turning yellow and stigmatic branches are starting to spread outwards (W9.5), we tag spikes to indicate the beginning of the time course. Depending on the location and scale of the experiment, logistical issues such as transport and preservation methods also need to be considered at the time of sampling. For sample collection, we carefully dissect wheat carpels from central spikelets in the field, which can be performed by eye as they are relatively large (Fig. 2B). Alternatively, we cut individual tillers between the uppermost and penultimate internode and transport them in water to the laboratory for carpel dissection. Once dissected, we store the carpels in 95% ethanol and acetic acid (75% v/v) for image acquisition at a later timepoint, or fresh (non-fixed) specimens are imaged directly if tillers have been transported to the laboratory. We use a stereo microscope with an integrated camera to acquire the two-dimensional RGB image of the carpel against a black background (Fig. 2C). We use different magnifications and fields of view to help capture the best representative plane of the carpel (Supplementary Figure 3). In the case of the fixed samples, we place carpels in a Petri dish with 70% ethanol to preserve the feathery appearance of the stigma. Only one image per sampled carpel is required for subsequent steps.

### Annotation of microscopy images and quantification of carpel traits

To process and perform quantitative analyses of the microscopy images, we trained a machine learning (ML)-based algorithm to automatically and rapidly annotate and measure stigma area and ovary diameter (Fig. 2D). The trained networks return a set of annotated images alongside their shape descriptors, together with a csv file containing the measurements of the analysed images in an output folder. This step requires some manual curation whereby the user inspects, detects, and corrects annotation errors, or removes corrupted images (Fig. 2E).

### Data modelling and knowledge transfer

We developed code to model growth dynamics of stigma area and ovary diameter (Fig. 2F). The open-access customizable R scripts can produce a range of outputs to compare genotypes, environmental conditions, or specific developmental stages, thereby helping to generate new hypotheses. Additionally, the exploitation of the knowledge generated could be key in the progress towards establishing successful hybrid breeding programmes as the selection of MS cultivars will be now based on novel phenotypic information.

### Development and validation of the stigma and ovary convolutional neural networks

Our aim was to develop an automated phenotyping method to detect and annotate the perimeter covered by the stigma hairs and the ovary to determine stigma area and ovary diameter across the life cycle of wheat carpels. To carry out automated image annotation and measurement, we trained a convolutional neural network (CNN) on a set of representative carpel images with manual annotations of the stigma perimeter (*n* = 86 images) and ovary perimeter (*n* = 121 images) using the UNet design [22] (Fig. 3A). The training dataset spanned a random sample of seven genotypes, ranging from early carpel development (W9.5 and earlier) to fully degenerated carpels (Fig. 1A), and included both fixed and non-fixed carpels. After successfully training the networks, we obtained an adapted stigma CNN able to quantify the area covered by stigmatic hairs and an adapted ovary CNN that, after some post-processing of the network output, quantifies the diameter of the ovary (Fig. 3B).

**Figure 3.**
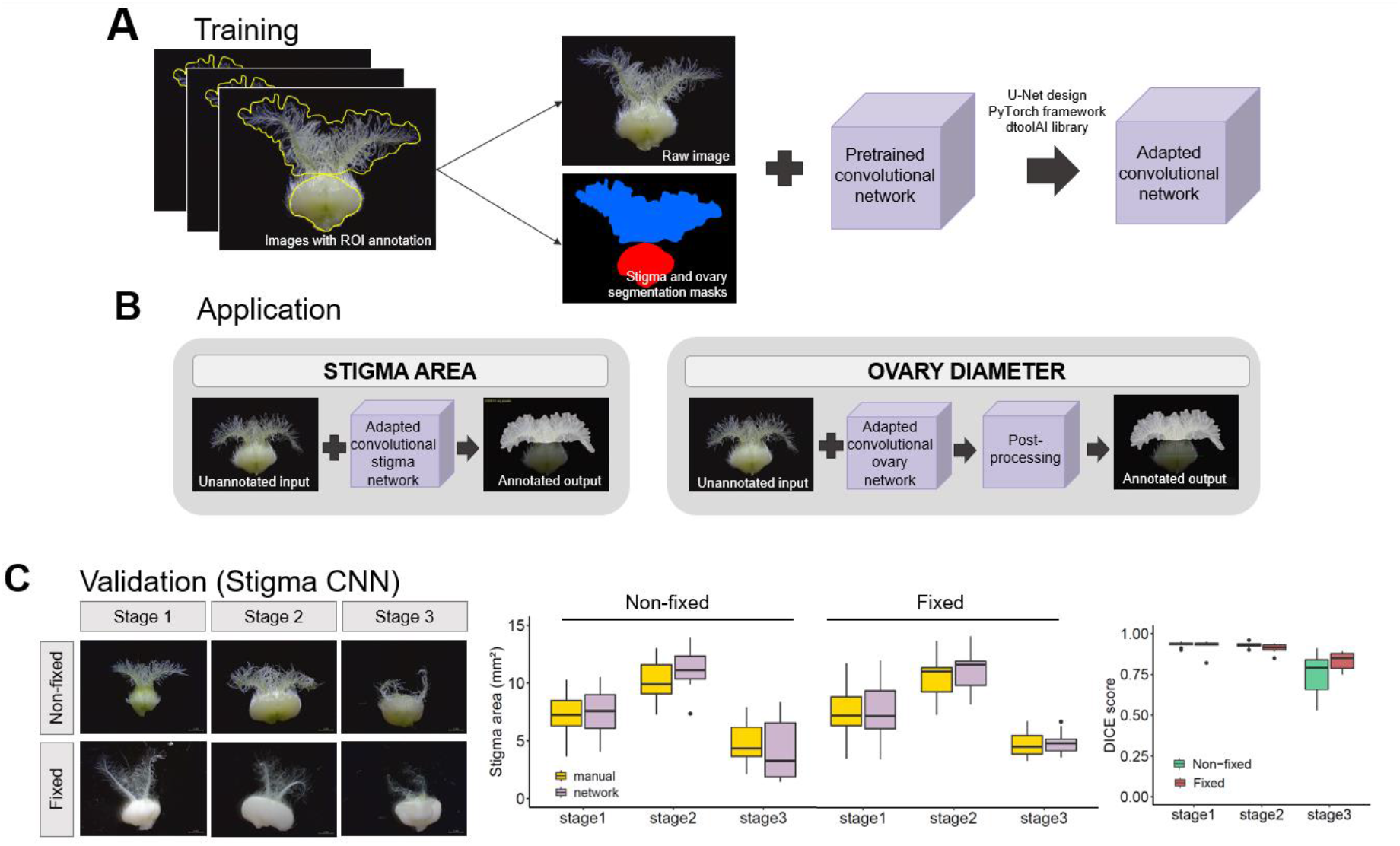
Development and validation of convolutional neural network for stigma and ovary annotation. **(A)** Pipeline for the development of the stigma and ovary networks. **(B)** Application of the adapted network for stigma area annotation to new data. **(C)** Cross-validation of ground-truth measurements and network values extracted from 60 randomly chosen images divided into six classes according to floral age (stage) and sampling method. First box plot shows the distribution of the stigma areas in mm^2^ per cross-validation class (*n* = 10 images per stage x method combination) determined by manual (yellow) and network (grey) annotation. There were no significant differences for any of the six classes (stage x sampling method combination). Second box plot indicates DICE scores of stigma area in non-fixed (green) and fixed (red) samples. A DICE score of 0 indicates no spatial overlap between the two sets of annotation results, 1 indicates complete overlap. The box plots show the middle 50% of the data with the median represented by the horizontal line. Whiskers represent datapoints within 1.5 times the interquartile range with outliers highlighted as individual points.

To evaluate the performance of the resulting adapted CNNs we applied the network to an unseen set of 60 microscope images without manual annotations (Fig. 3C). Subsequently, we manually annotated this new dataset using Fiji to provide a ground truth reference with which to compare to the CNN annotated outputs. We divided the cross-validation process according to the developmental stage of the carpels, sampling method (fixed or non-fixed), and the tissue of interest (Fig. 3C and Supplementary Figure 4 for ovary CNN validation). We observed a high overlap in stigma areas and ovary diameters of fixed and non-fixed samples between the manual and automated annotations across all three developmental stages. There were no significant differences between the manual and automated annotations, apart for ovary diameter of the non-fixed samples at stage 3 (*P* = 0.03, one-way ANOVA; Supplementary Table 1). Additionally, we calculated the Dice Similarity Coefficient (DSC) for each group of images which allowed quantitative evaluation of the performance of the adapted networks (Fig. 3C and Supplementary Figure 4). Overall, we found very uniform DSC values between the ground-truth and CNN annotation across floral traits, sampling methods and developmental stages (with the exception of the stigma CNN at the last developmental stage) with DSC mean averages of 0.89 and 0.95 for the stigma and ovary CNN, respectively. Together, these results show that our machine learning approach quantifies key parameters of the carpel life cycle in wheat in agreement with the more time-consuming manual measurements.

### Variation in stigma and ovary growth patterns can be studied on fixed carpels

Chemical fixation is commonly used to prevent tissue autolysis and degradation, while preserving morphology and cellular details for subsequent macro or microscopic evaluations. Fixatives, however, can lead to changes in volume and shape of the treated specimens due to cell shrinkage or swelling. Thus, artefacts of the technique could potentially lead to erroneous conclusions when measuring morphological traits of fixed samples.

To assess the effect of the fixative solution (ethanol and acetic acid) on stigma area and ovary diameter, we used four MS cultivars and sampled carpels at five timepoints over an 18-days period. Analysis of variance (three-way ANOVA) indicated that the fixative significantly reduces stigma area (*P* < 0.001), whereas ovary diameter remained unchanged after applying the fixative (*P* = 0.25) (Fig. 4A and Supplementary Figure 5). For stigma area, the fixative x timepoint interaction was borderline non-significant (*P* = 0.09) (Supplementary Table 2) suggesting that the response to the fixative might change with floral age. By analysing individual timepoints, we observed that at 3 and 7 days after W9.5, at the peak of the stigma area, there are no-significant differences between fixed and non-fixed samples, whereas at 0, 13 (*P* < 0.05) and 18 days after W9.5 (*P* < 0.001) (Fig. 4A) the fixed samples show a reduced stigma area. Importantly, the absence of a significant fixative x cultivar interaction (*P* > 0.52) suggests that all four cultivars react to the fixative in a similar manner (Fig. 4D). Taken together, we see that fixing the carpels in ethanol and acetic acid reduces stigma area although the developmental dynamics of stigma area are conserved in the four cultivars across the 18 days (Fig. 4D). We therefore conclude that the use of ethanol and acetic acid fixative allows us to accurately capture the growth dynamics of stigma area and ovary diameter and investigate phenotypic variation among diverse genotypes.

**Figure 4.**
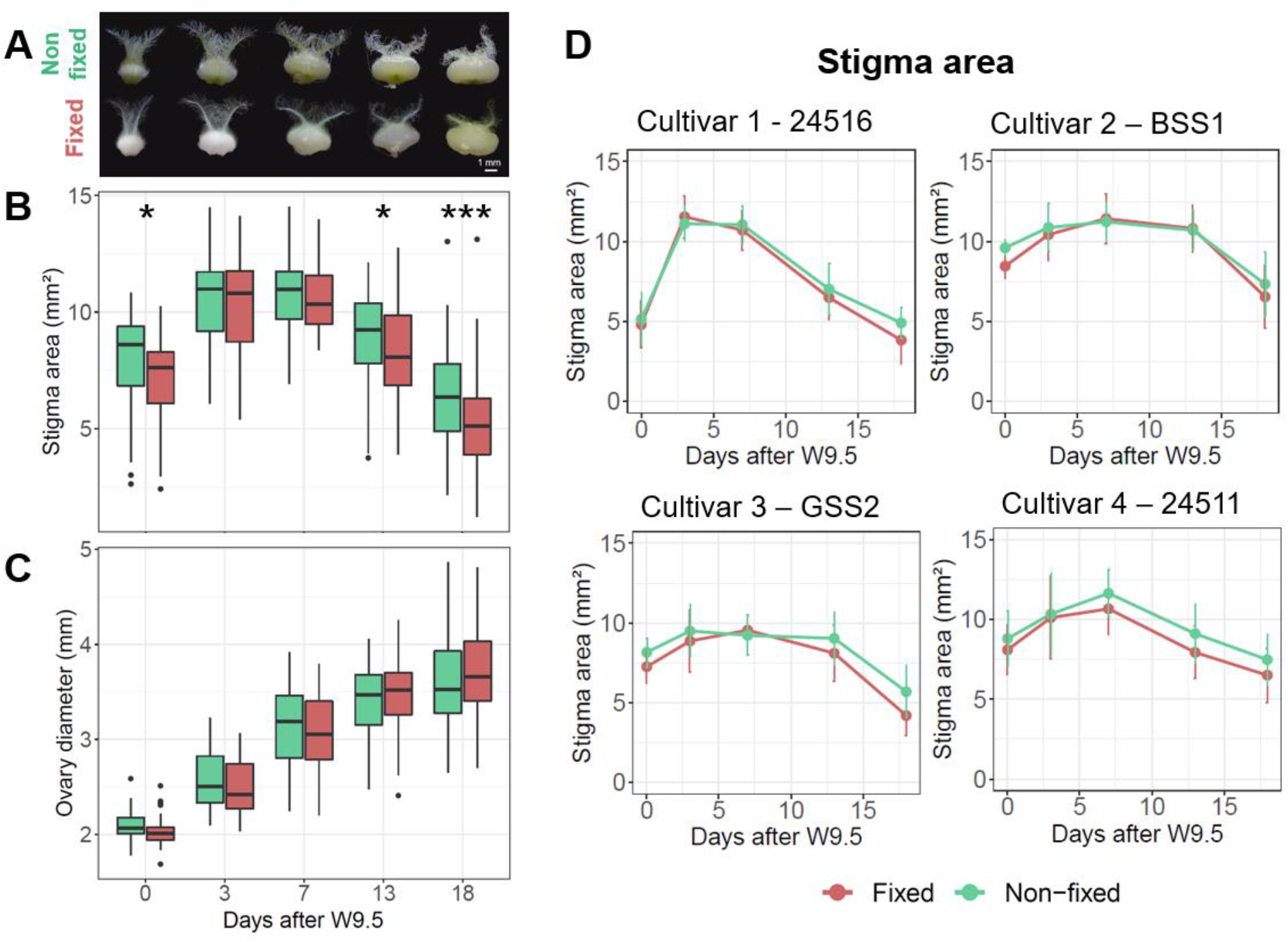
Effects of the fixative on carpel morphology across time and cultivars. **(A)** Representative images of carpels before (non-fixed) and after (fixed) applying the fixative. **(B-C)** Box plots showing the comparison between non-fixed (green) and fixed (red) samples for stigma area **(B)** and ovary diameter **(C)** at different sampling points. Data are the average of the four cultivars shown in panel D which comprise 10-20 carpels from a total of 4 plants per timepoint and cultivar. * *P* <0.05; \*\*\**P* <0.001. The box plots show the middle 50% of the data with the median represented by the horizontal line. Whiskers represent datapoint within 1.5 times the interquartile range with outliers highlighted as individual. **(D)** Developmental dynamics of stigma area of four male sterile cultivars, comparing non-fixed (green) and fixed (red) carpel samples (between 10-20 carpels from a total of 4 plants per timepoint). Error bar denotes the standard error.

Nonetheless, caution must be taken to compare absolute stigma areas across development given the significant reduction at early and late timepoints.

### The application of the phenotyping approach provides insight into the developmental behaviour of the unpollinated wheat carpel

Having established the method to quantitively measure the progression of carpel development in the absence of pollination, we next sought to employ this approach to gain insights into genetic and environmental variation for these two floral traits. To accomplish this, we applied our phenotyping approach to three MS cultivars grown during two consecutive field seasons (2020 and 2021) where we performed a developmental time course ranging from W9.5 until the carpel had visually deteriorated. To accommodate for season-specific differences in temperature between the two seasons (Supplementary Figure 2), we incorporated daily temperatures in our model to normalise developmental stages by cumulative degree days.

We found that all three MS cultivars exhibit contrasting developmental patterns for stigma area and ovary diameter and that these differences among cultivars are largely maintained across field seasons (Fig. 5A,B). The phenotypic differences, particularly in stigma area, are observed in both the growth (positive slope) and deterioration (negative slope) phases of carpel development, which inevitably impacts on the overall duration of the life cycle. For instance, we can distinguish the fast development of carpels from cultivar 24522 from the slow progression of carpels from cultivar BSS1 (Fig. 5A). Despite these differences, all three patterns seem to underline a common developmental trend for the dynamics of stigma and ovary traits that is characterised by: (1) an initial growth phase, (2) followed by a peak phase in which stigma reaches its maximum and the radial expansion of the ovary slows down, and (3) a final deterioration phase. This conceptual framework for quantifying and classifying the development of the unpollinated carpel is presented in Fig. 5C,D. The results obtained from breaking down late carpel development into more descriptive phases are detailed below.

**Figure 5.**
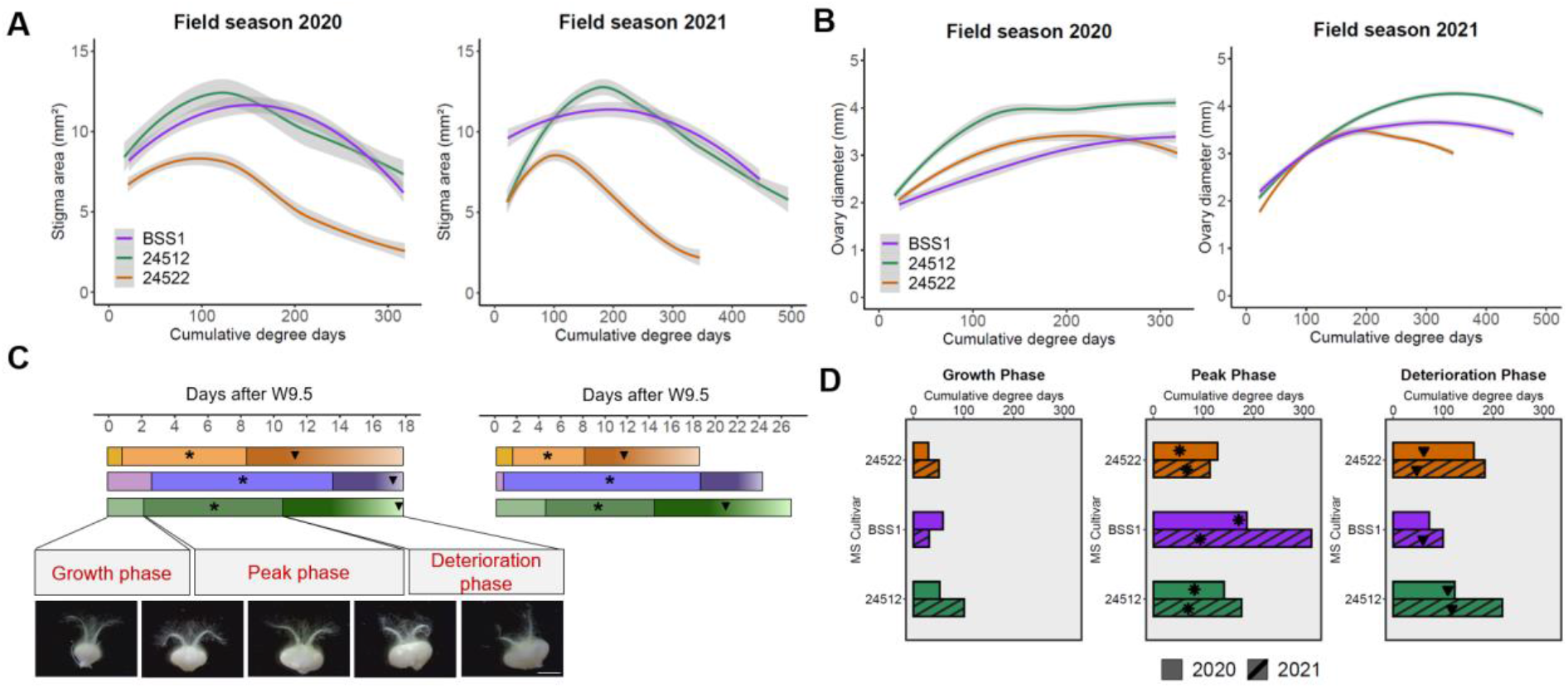
Phenotypic quantification of carpel development in three distinct MS cultivars under field conditions. **(A)** Temporal trends of stigma area (mm^2^) for 2020 and 2021 field seasons. **(B)** Temporal trends of ovary diameter (mm) in 2020 and 2021. **(C)** Growth, peak and deterioration phases represented in days after W9.5 (temporal units are directly comparable to cumulative degree days in panel A). Representative images illustrating carpel appearance at the beginning and end of each phase (3^rd^ picture represent carpel with 100% stigma area). **(D)** Bar charts show the duration (in degree days) of each of the phases across cultivars and field seasons. The end of the deterioration phase is marked by the last sampling point. In A and B, polynomial regression models at a 95% confidence interval (Loess smooth line) are shown. Grey shading represents standard error. Five carpels from 4 and 6 plants were sampled at each timepoint in 2020 and 2021, respectively. Scale bar in C = 2 mm. C and D, (*) indicates maximum stigma area; (▼) indicates a 40% drop in stigma area with respect to the maximum area. Before plotting, outliers were filtered out following the interquartile range criterion.

### Growth phase

Stigma and ovaries experience rapid and exponential growth during the first phase. The growth phase is underway at W9.5 (around ear emergence) and extends for 1 to 4 days until the stigma is well developed and potentially receptive for pollination. The end of the phase coincides with a developmental stage parallel to W10 (anthesis) in male-fertile plants. We found that for all three cultivars the end of this phase could be described by the stigma showing an area of approximately 85% of its maximum size. The criteria for us to select the 85% cut-off as the end of the growth phase was based on 85% being the percentage that was present in all three cultivars across both field seasons and happened shortly after W9.5, thus mimicking anthesis (Supplementary Figure 6).

### Peak phase

This second phase is denoted by the stigmatic tissue reaching its maximum size (asterisks in Fig. 5C) at around 5 to 10 days after W9.5 (depending on the cultivar and year). After reaching this peak, a gradual and irreversible decline in stigma area is accompanied by a notable arrest of the ovary radial expansion. To mirror the behaviour of stigma area at the beginning of this phase, we selected a 15% drop in stigma area to mark the end of the peak phase. Using this classification, we observed that this phase extends until 8 to 14 days after W9.5 in 2020, and 8 to 18 days in 2021 (Fig. 5C). Considering previous studies on female receptivity in wheat [28-30] we hypothesise that the wheat carpel shows its maximum reproductive potential during this phase.

### Deterioration phase

During this phase, symptoms of stigma deterioration start to become obvious, where clusters of stigma hairs are collapsing in response to a loss in turgor pressure (inverted triangles in Fig. 5C indicate a 40% drop in stigma area). The collapse of the remaining stigma hairs continues for several days resulting in a completely deteriorated stigma at 18 and 27 days after W9.5 in 2020 and 2021, respectively (Fig. 5C,D). By the end of this phase the ovary walls also show an irregular surface due to tissue deterioration. Based on these observations, we speculate that the onset of this phase marks the irrevocable loss of the reproductive potential of the floret.

Dissecting carpel development into growth, peak and deterioration phases allowed us to assign the cultivars BSS1, 24512 and 24522 into slow, moderate, and fast developing carpels, respectively, according to when they reach the beginning of the deterioration phase. For example, cultivar BSS1 reached the onset of the deterioration phase at 14 and 18 days after W9.5 in 2020 and 2021, respectively. This was approximately 4 days after cultivar 24512 and between 6 to 10 days after cultivar 24522 in 2020 and 2021, respectively (Fig. 5C). The developmental pattern classification also allows comparisons across field seasons, where the relative ranking of cultivars was well conserved between years. Although we adjusted for cumulative degree days, the colder and damper weather conditions of 2021 were reflected in an extension in the duration of most phases in some, but not all cultivars (Fig. 5D). For example, duration of the three phases was largely unaffected in cultivar 24522, whereas in cultivar BSS1, the duration of the peak phase in 2021 (315 degree days) was almost twice that of the previous year (186 degree days). These initial analyses suggest that while the developmental dynamics of stigma area and ovary diameter are largely consistent across years there are cultivars in which the duration of the developmental phases could be sensitive to other environmental changes that remain to be tested.

## Discussion

### High throughput phenotyping for the quantification of floral traits in unpollinated wheat carpels

Our understanding of floral developmental processes has been assisted by the establishment of scales that describe changes in the shape, size and surface features of floral organs leading up to anthesis [31, 32]. These scales often divide a continuous developmental process into defined stages, which are characterised by landmark events, such as the appearance of stigmatic branches. These scales have facilitated the interpretation of genetic studies and have contributed towards our understanding of the mechanisms that underlie the transitions leading to a given landmark event. During post-pollination stages, the focus shifts towards the developing fruit. Previous work, however, has also illustrated the importance of the quantitative monitoring of morphological changes associated with late carpel development (i.e., in the absence of pollination), such as silk elongation in maize [9] or ovary radial expansion in wheat [11], as they represent survival mechanisms to ensure seed set by cross-pollination. So far, the few studies investigating the progression of the unpollinated female carpel after anthesis have focused in giving detailed descriptions of flowers from one or two different genotypes grown in controlled environment conditions. These types of meticulous approaches are arduous and expensive not only to implement in large scale experiments but also to execute under field conditions, where equipment is often limited. Consequently, studies investigating detailed phenotypes in the field are lacking.

To enhance our understanding of the biological processes that occur in unpollinated carpel under breeding-relevant conditions, we created a machine learning-based approach to phenotype field-grown MS wheat cultivars. Given the sequence of morphological changes we observed in the unpollinated wheat carpel (Fig. 1A), we next quantified changes in stigma area and ovary diameter to describe carpel development. Our findings are two-fold: (1) we demonstrate the suitability of our approach for the detailed study of floral organ traits in field screenings, and (2) we show that the unpollinated carpel undergoes a well-defined pattern of growth and senescence characterised by gradual changes in stigma and ovary sizes (Fig. 5A,B). Based on these findings, we propose developmental phases that are relative to the maximum stigma area and ovary size (Fig. 1C and 5C) with which to build the foundations of future research of floral organ development and senescence in the absence of pollination.

### Considerations on the use of the stigma and ovary adapted CNNs

The quantification of stigma area and ovary diameter cannot be easily determined from surface observations of wheat spikes and requires the dissection and microscopy of individual carpels. The manual annotation and quantification of microscopy images is cumbersome and often delays scientific discoveries. Deep learning-based approaches, such as Convolutional Neural Networks (CNNs), have emerged as a solution to perform image quantification in an automated, rapid, and less biased manner, lifting the burden of image analysis from researchers. In this work we developed two CNNs (both publicly available at https://github.com/marina-millan/ML-carpel_traits) that enable non-machine-learning experts to quantify stigma area and ovary diameter on their local computer in a matter of hours (Fig. 3). Evaluation metrics on the performance of the adapted networks (Fig. 3C and Supplementary Table 1) demonstrate their capability to satisfactorily measure both floral traits by condensing each RGB image to a single value (i.e., pixels).

Measurements between manual and CNN annotation were largely indistinguishable (Supplementary Table 1), with the network being less capable at later stages (Fig. 3C, Supplementary Figure 1). We believe one of the reasons for the poorer performance could be due to the difficulty in distinguishing the stigma and ovary from each other when the stigma is severely deteriorated and resting on top of the ovary (Supplementary Figure 7). Also, it is worth noting that the ovary CNN relies on the performance of the stigma CNN to correctly predict the diameter of the ovary (see Materials and Methods for the detailed description of the algorithm), such that a bad prediction of the stigma area will likely affect ovary annotation. Poor quality images (i.e., out of focus, poor resolution images) and certain carpel orientations (Supplementary Figure 3) also hinder the identification of the ovary and/or stigma, impairing the normal performance of the CNNs. In the case of blurry images, using post-processing tools (e.g., Photoshop or other image sharpening techniques) to adjust the sharpness of the image might help reduce the likelihood of incorrect annotations. However, there will still be certain cases where there is not an immediate reason for the failure of the CNN. We suggest, therefore, including an additional output verification step (Fig. 2E) to identify potential errors before continuing with the downstream analyses.

### The implementation of the phenotyping approach opens new research paths on the biology of late carpel development

To gain a more comprehensive overview of the developmental dynamics of the unpollinated carpel, we used our phenotyping approach (Fig. 2) to examine the sequential progression of changes in stigma and ovary morphology in three MS cultivars over two field seasons (Fig. 5). Across cultivars and seasons, we were able to identify an initial stigma and ovary growth phase, followed by a peak phase describing carpel developmental maturity, and a subsequent deterioration phase characterised by the eventual collapse of the female reproductive tissues (Fig. 5C). Equivalent patterns for the post-anthesis development of the unpollinated stigma have also been reported in maize, peas (*Pisum sativum*) and *Arabidopsis* [9, 14, 33], suggesting a conserved developmental programme that ensues in the absence of pollination.

Despite the conserved overall patterns, we identified differences in the duration of the growth, peak and deterioration phases in the three wheat MS cultivars used here. Gene expression studies of unpollinated *Arabidopsis* carpels [14] and stigma [15] have demonstrated that the lifespans of these floral structures are controlled by transcription factors that regulate developmental programmed cell death in these tissues. The results from *Arabidopsis* raise the prospect that the phenotypic variation observed between the three wheat MS cultivars could be due to differential gene expression patterns across cultivars that alter the onset of stigma senescence (Fig. 5A,C). Thus, new transcriptomic studies investigating the developmental transitions observed among the different field-grown cultivars would contribute to our understanding of the mechanisms governing these phases. We also observed that MS cultivars 24512 and 24522 had largely equivalent peak phase durations across years, whereas the duration of the peak phase in the CMS cultivar BSS1 varied almost two-fold between field seasons (Fig. 5D). This suggests that, despite accounting for temperature in our analyses (by using cumulative degree days), the duration of the stigma peak phase is sensitive to additional environmental factors. The response to these additional environmental factors could depend on the genotype, sterility system used, and/or the developmental phase in which the environmental stimuli are encountered. Consistent with this, several studies in wheat have attributed differential seed set rates of out-crossing MS plants (i.e., an indicator for the duration of stigma receptivity) to environmental factors such as temperature, relative humidity, and soil water availability [5, 30, 34]. Therefore, additional studies under field and controlled environment conditions will shed light on the causalities for the variation observed in ovary and stigma development across field seasons. Our phenotyping approach now improves the accessibility of the wheat carpel to detailed phenotypic analyses of the size of populations that are used in breeding programmes. This facilitates the identification of mutations that underpin genetic variation in carpel development contributing to understand gene function on a genome-wide scale. All in all, we provide a framework in which to conduct these new studies targeting diverse environments and genotypes, facilitating future hypothesis generation not only in wheat but also in other cereal crops.

### First steps towards an integrated developmental scale of the unpollinated wheat flower

The ultimate role of the carpel is the production of a viable seed. Thus, increasing the functional lifespan of carpels and stigmas (i.e., floral receptivity) are desirable agronomic traits that have the potential to increase the effective pollination period and seed set [35]. Yet, detailed evaluations of carpel and stigma development and how they relate to female floral receptivity and seed set are still lacking, even more so in cereals. It is reasonable to think that the functional lifespan of stigma receptivity would coincide with stigma cell integrity, as illustrated in early studies of kiwifruit (*Actinidia deliciosa*) and maize [9, 16]. According to these studies we could, for example, speculate that (a) seed set rates will be higher if pollination occurs during the peak phase compared to the deterioration phase, or (b) that cultivars with a prolonged peak phase (such as BSS1) will be receptive to pollination for longer than cultivars with a shorter peak phase (such as 24512 or 24522) (Fig. 5C). However, as recently demonstrated in *Arabidopsis*, a delay in stigma senescence caused by the disruption of two programmed cell death-promoting transcription factors was only accompanied by a minor extension in floral receptivity, suggesting that additional processes must be involved in controlling the duration of floral receptivity [15]. New studies, therefore, need to be conducted to help investigate stigma receptivity under defined phases of carpel development to help clarify the relationship between stigma morphology and viability, pollen germination, and seed set. Additionally, such information will allow a greater understanding of how genetic and environmental factors affect various aspects of the stigma life cycle (e.g., loss of stigma receptivity, onset of stigma cell death).

The next steps towards understanding the cross-pollination process in the field will also require integrating the changes in carpel morphology with those of the overall spike. For instance, as reviewed by Selva et al. [19], certain wheat spike architectures, like the openness of the floret, facilitate airborne pollen access which would additionally contribute to increasing out-crossing rates in hybrid production.

Our approach for phenotyping carpel development provides a new tool for examining a fertility trait that is poorly understood and hitherto time-consuming to analyse. Together with recent advances in genetic resources [36-38] and genome sequence data [39, 40], this approach provides a new opportunity to unlock genetic variation for stigma and ovary traits that associate with floret fertility, which is vital given that improved fertilisation will help address the increasing demands to enhance global food production.

## Supplementary Table Captions

**Supplementary Table 1**. Summary table on the manual and CNN annotation metrics obtained for stigma area and ovary diameter. Estimated marginal means ± standard errors are given for each type of annotation. P values indicate statistical significance of the pairwise comparison in one-way ANOVA.

**Supplementary Table 2**. Summary table of three-way ANOVA on the effect of the fixative on stigma area and ovary diameter across different timepoints. P values shown for the different timepoints indicate the P values for the pairwise comparisons. Estimated marginal means ± standard errors are also given for each treatment (fixed or non-fixed).

## Supplementary Figure Captions

**Figure S1**. Schematic representations of the field layout.

**Figure S2**. Environmental conditions recorded during 2020 and 2021 field seasons.

**Figure S3**. Unaccepted and accepted carpel images.

**Figure S4**. Validation of convolutional neural network for stigma and ovary annotation.

**Figure S5**. Effects of the fixative on ovary diameter across time and cultivars.

**Figure S6**. Developmental stigma patterns expressed in percentage from maximum observed stigma area.

**Figure S7**. Erroneous stigma area and ovary diameter CNN annotations.

## Author contributions

Contributions according to CRediT (https://casrai.org/credit/): Conceptualization: MMB, CU, SB; Data curation: MMB; Formal Analysis: MMB; Funding acquisition: SB; Investigation: MMB; Methodology: MMB; Software: MH; Supervision: SB, CU; Visualization: MMB, MH, CU; Writing – original draft: MMB, MH, SB, CU; Writing – review & editing: all authors

## Funding

This work was supported by the UK Biotechnology and Biological Sciences Research Council (BBSRC) through the Designing Future Wheat (BB/P016855/1) and Genes in the Environment (BB/P013511/1) Institute Strategic Programmes. Additional funding was provided by the European Research Council (ERC-2019-COG-866328). MMB was supported by a BBSRC Norwich Research Park Biosciences Doctoral Training Grant (BB/M011216/1).

## Competing Interest

NB is employed by KWS UK Ltd and YM is employed by Syngenta, France. The remaining authors declare that they have no conflict of interest.

## Data Availability Statement

For enquiries regarding the germplasm used in this work contact NB or YM. Training codes used for the development of the CNNs, adapted stigma and ovary CNNs and R scripts used for data curation and visualisation can be found here: https://github.com/marina-millan/ML-carpel_traits

## Acknowledgment

We thank the JIC Field Trials and Horticultural Services teams for technical support in field and glasshouse experiments, the JIC Bioimaging facility and staff for their contribution to this publication and Professor Lars Østergaard (JIC) and Dr. Azahara Martín for feedback on the work. We also want to thank James Simmonds and Tobin Florio for their assistance during the field trials and Flozbox studio (https://flozbox-science.com/) for designing Fig. 2. For the purpose of open access, the author has applied a Creative Commons Attribution (CC BY) licence to any Author Accepted Manuscript version arising.

## References

1. Walker, E.R., On the Structure of the Pistils of Some Grasses University studies from University of Nebraska, 1906. Vol. VI (No.3).

2. Heslop-Harrison, Y. and K. Shivanna, The receptive surface of the angiosperm stigma. Annals of botany, 1977. 41(6): p. 1233–1258.

3. Heslop-Harrison, J., Pollen-stigma interaction in grasses: a brief review. New Zealand Journal of Botany, 1979. 17(4): p. 537–546.

4. Edlund, A.F., R. Swanson, and D. Preuss, Pollen and stigma structure and function: the role of diversity in pollination. Plant Cell, 2004. 16 Suppl: p. S84–97.

5. Fabian, A., et al., Stigma Functionality and Fertility Are Reduced by Heat and Drought Co-stress in Wheat. Front Plant Sci, 2019. 10: p. 244.

6. Onyemaobi, I., et al., Both Male and Female Malfunction Contributes to Yield Reduction under Water Stress during Meiosis in Bread Wheat. Front Plant Sci, 2016. 7: p. 2071.

7. Mitchell, J.C. and J.F. Petolino, Heat Stress Effects on Isolated Reproductive Organs of Maize. Journal of Plant Physiology, 1988. 133(5): p. 625–628.

8. Kempe, K. and M. Gils, Pollination control technologies for hybrid breeding. Molecular breeding, 2011. 27(4): p. 417–437.

9. Westgate, P.B.a.M.E., Emergence, Elongation, and Senescence of Maize Silks. Crop Science, 1993. 33: p. 271–275.

10. Molnár-Láng, M., B. Barnabas, and E. Rajki, Changes in the shape, volume, weight and tissue structure of the pistil in the flowers of male sterile wheats during flowering. Cereal Research Communications, 1980: p. 371–379.

11. Okada, T., et al., Unfertilized ovary pushes wheat flower open for cross-pollination. J Exp Bot, 2018. 69(3): p. 399–412.

12. Primack, R.B., Longevity of individual flowers. Annual review of ecology and systematics, 1985. 16(1): p. 15–37.

13. Ashman, T.-L. and D.J. Schoen, How long should flowers live? Nature, 1994. 371(6500): p. 788–791.

14. Carbonell-Bejerano, P., et al., A fertilization-independent developmental program triggers partial fruit development and senescence processes in pistils of Arabidopsis. Plant Physiol, 2010. 154(1): p. 163–72.

15. Gao, Z., et al., KIRA1 and ORESARA1 terminate flower receptivity by promoting cell death in the stigma of Arabidopsis. Nature plants, 2018. 4(6): p. 365–375.

16. González, M.V., M. Coque, and M. Herrero, Papillar integrity as an indicator of stigmatic receptivity in kiwifruit (Actinidia deliciosa). Journal of Experimental Botany, 1995. 46(2): p. 263–269.

17. Pieruschka, R. and U. Schurr, Plant Phenotyping: Past, Present, and Future. Plant Phenomics, 2019. 2019: p. 1–6.

18. Furbank, R.T. and M. Tester, Phenomics – technologies to relieve the phenotyping bottleneck. Trends in Plant Science, 2011. 16(12): p. 635–644.

19. Selva, C., et al., Hybrid breeding in wheat: how shaping floral biology can offer new perspectives. Funct Plant Biol, 2020. 47(8): p. 675–694.

20. Waddington, S., P. Cartwright, and P. Wall, A quantitative scale of spike initial and pistil development in barley and wheat. Annals of Botany, 1983. 51(1): p. 119–130.

21. Zadoks, J.C., T.T. Chang, and C.F. Konzak, A decimal code for the growth stages of cereals. Weed research, 1974. 14(6): p. 415–421.

22. Falk, T., et al., U-Net: deep learning for cell counting, detection, and morphometry. Nature methods, 2019. 16(1): p. 67–70.

23. Adam Paszke, S.G., Francisco Massa, Adam Lerer, James Bradbury, Gregory Chanan, Trevor Killeen, Zeming Lin, Natalia Gimelshein, Luca Antiga, Alban Desmaison, Andreas Kopf, Edward Yang, Zachary DeVito, Martin Raison, Alykhan Tejani, Sasank Chilamkurthy, Benoit Steiner, Lu Fang, Junjie Bai, Soumith Chintala, PyTorch: An Imperative Style, High-Performance Deep Learning Library, in 33rd Conference on Neural Information Processing Systems. 2019, NeurIPS Proceedings: Vancouver.

24. Hartley, M. and T.S.G. Olsson, dtoolAI: Reproducibility for Deep Learning. Patterns, 2020. 1(5): p. 100073.

25. Dice, L.R., Measures of the Amount of Ecologic Association Between Species. Ecology, 1945. 26(3): p. 297–302.

26. Diederik P. Kingma, J.L.B., ADAM: A METHOD FORSTOCHASTICOPTIMIZATION, in 3rd International Conference for Learning Representations. 2015: San Diego.

27. Perry Miller, W.L., Stu Brandt, Using Growing Degree Days to Predict Plant Stages. Montguide, 2001. MT200103 AG 7/2001.

28. Pickett, A.A., Hybrid wheat-results and problems. Fortschritte der Pflanzenzüchtung (Germany), 1993.

29. Kirby, E., Botany of the wheat plant. Bread Wheat. Improvement and Production. Food and Agriculture Organization of the United Nation. Rome, 2002: p. 19–37.

30. De Vries, A.P., Flowering biology of wheat, particularly in view of hybrid seed production—a review. Euphytica, 1971. 20(2): p. 152–170.

31. S.R. Waddington, P.M.C., A Quantitative Scale of Spike Initial and Pistil Development in Barley and Wheat. 1983.

32. David R. Smyth, J.L.B., and Elliot M. Meyerowitz, Early Flower Development in Arabidopsis. The plant cell, 1990.

33. Y. Vercher, A.M. C. López, J.L. García-Martínez, J. Carbonell, Structural changes in the ovary of Pisum sativum L. induced by pollination and gibberellic acid. Plant Science Letters, 1984. 36(2): p. 87–91.

34. Imrie, B., Stigma receptivity in cytoplasmic male sterile wheat. Australian Journal of Experimental Agriculture, 1966. 6(21): p. 175–178.

35. Williams, R., The effect of summer nitrogen applications on the quality of apple blossom. Journal of Horticultural Science, 1965. 40(1): p. 31–41.

36. Krasileva, K.V., et al., Uncovering hidden variation in polyploid wheat. Proceedings of the National Academy of Sciences, 2017. 114(6): p. E913–E921.

37. Wingen, L.U., et al., Establishing the A. E. Watkins landrace cultivar collection as a resource for systematic gene discovery in bread wheat. Theoretical and Applied Genetics, 2014. 127(8): p. 1831–1842.

38. Sansaloni, C., et al., Diversity analysis of 80,000 wheat accessions reveals consequences and opportunities of selection footprints. Nature communications, 2020. 11(1): p. 1–12.

39. Walkowiak, S., et al., Multiple wheat genomes reveal global variation in modern breeding. Nature, 2020. 588(7837): p. 277–283.

40. International Wheat Genome Sequencing, C., et al., Shifting the limits in wheat research and breeding using a fully annotated reference genome. Science, 2018. 361(6403).

