## Supplementary Figures for "A scalable phenotyping approach for female floral organ development and senescence in the absence of pollination in wheat"

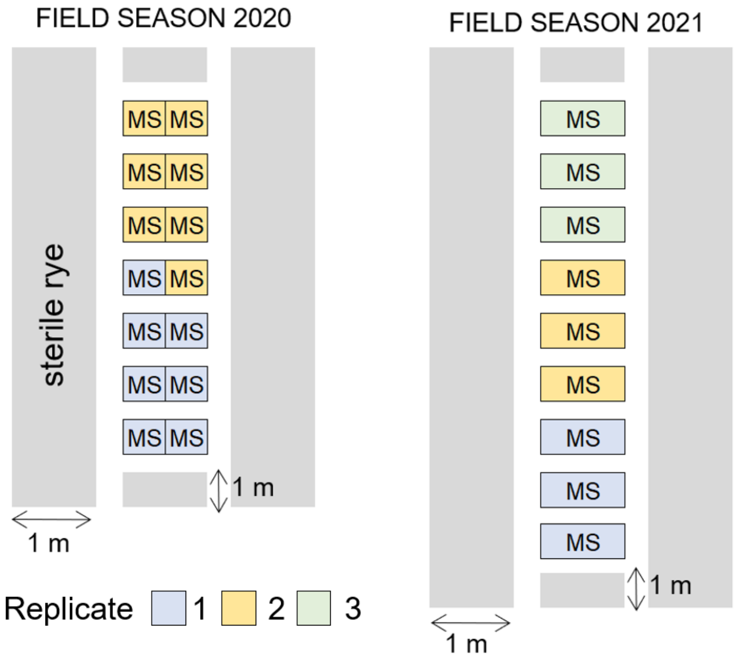


**Figure S1. Schematic representations of the field layout.** Male sterile (MS) cultivars were grown surrounded by a continuous stripe of sterile rye that was used as pollen barrier. Plots were replicated twice in 2020 (*N* = 7) and 3 times in 2021 (*N* = 3).

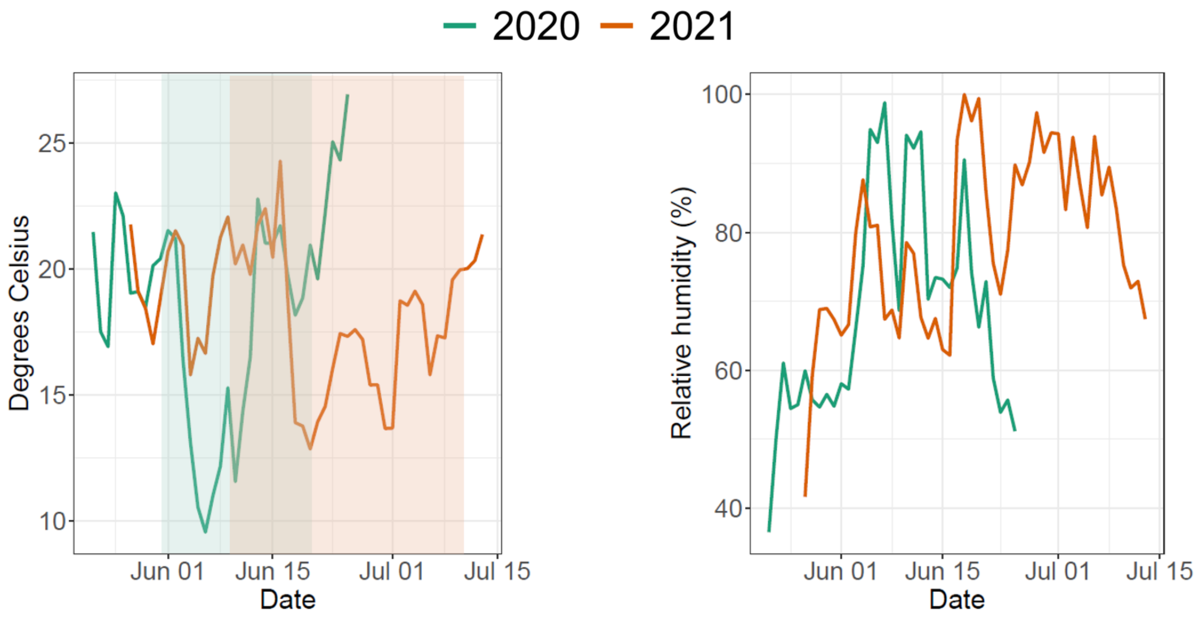


**Figure S2. Environmental conditions recorded during 2020 and 2021 field seasons.** Left panel illustrates daily temperatures in degrees Celsius recorded during both field experiments (2020: green line; 2021: orange line). Shaded rectangles indicate the beginning and end of 2020 and 2021 time courses. Right panel shows the water vapor contained in the air expressed in percentage of relative humidity.

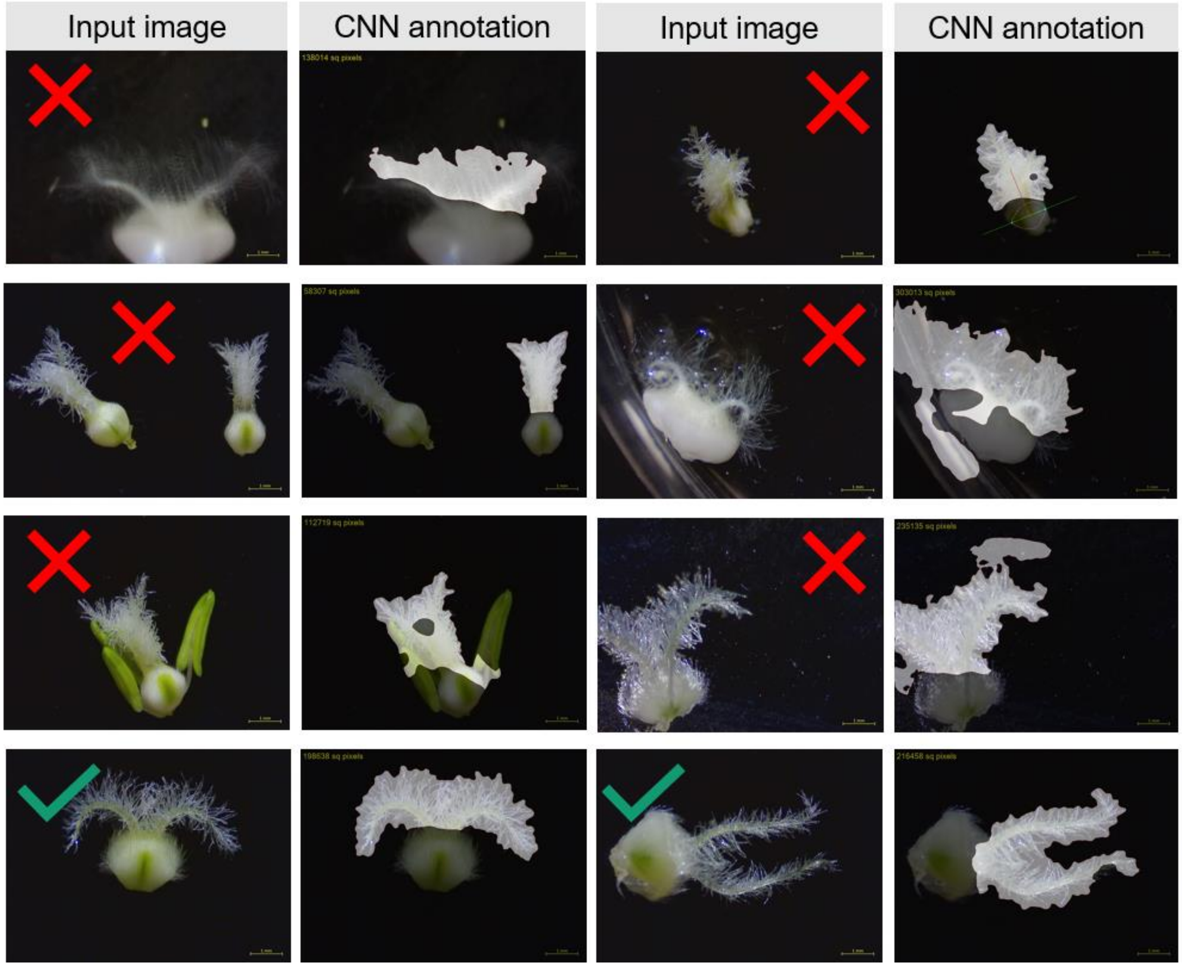


**Figure S3. Unaccepted and accepted carpel images.** Representation of images (jpeg format) unlikely to be correctly annotated by either the stigma CNN or ovary CNN (red crosses, e.g., blurry or out of focus images, lifted carpels, presence of other objects, etc.) and example of images the stigma and ovary CNNs expect and would be able to annotate accurately (green ticks).

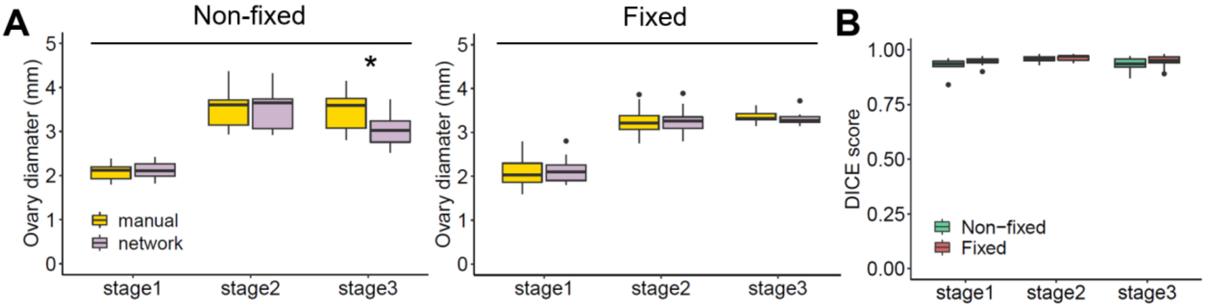


**Figure S4. Validation of convolutional neural network for stigma and ovary annotation**. **(A)** Cross-validation of ground-truth measurements and network values extracted from 60 randomly chosen images divided into six classes according to floral age and sampling method. Distribution of the ovary diameter in mm per cross-validation class (*n* = 7-10), determined by manual (yellow) and automated (violet) annotation. **(B)** Box plots showing Dice similarity coefficient of ovary diameter in non-fixed (green; *n* = 10) and fixed (red; *n* = 10) samples (0 indicates no spatial overlap between the two sets of annotation results, 1 indicates complete overlap). Box plots show the middle 50% of the data with the median represented by the horizontal line. Whisker represents datapoint within 1.5 times the interquartile range with outliers highlighted as individual.

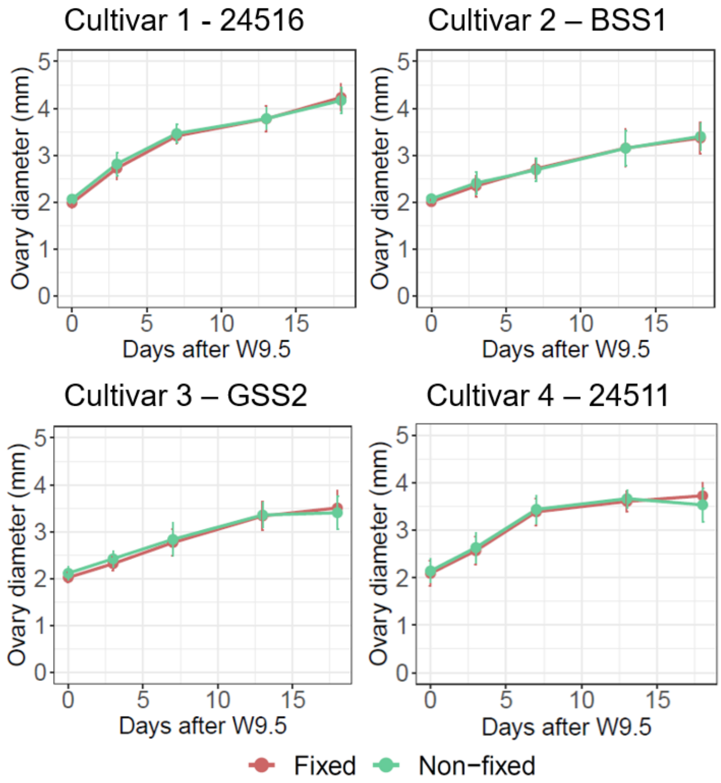


**Figure S5. Effects of the fixative on ovary diameter across time and cultivars.** Stigma area development dynamics of four MS cultivars comparing non-fixed (green) and fixed (red) carpel samples (between 10-20 carpels from 4 plants per timepoint). Error bar denotes the standard error.

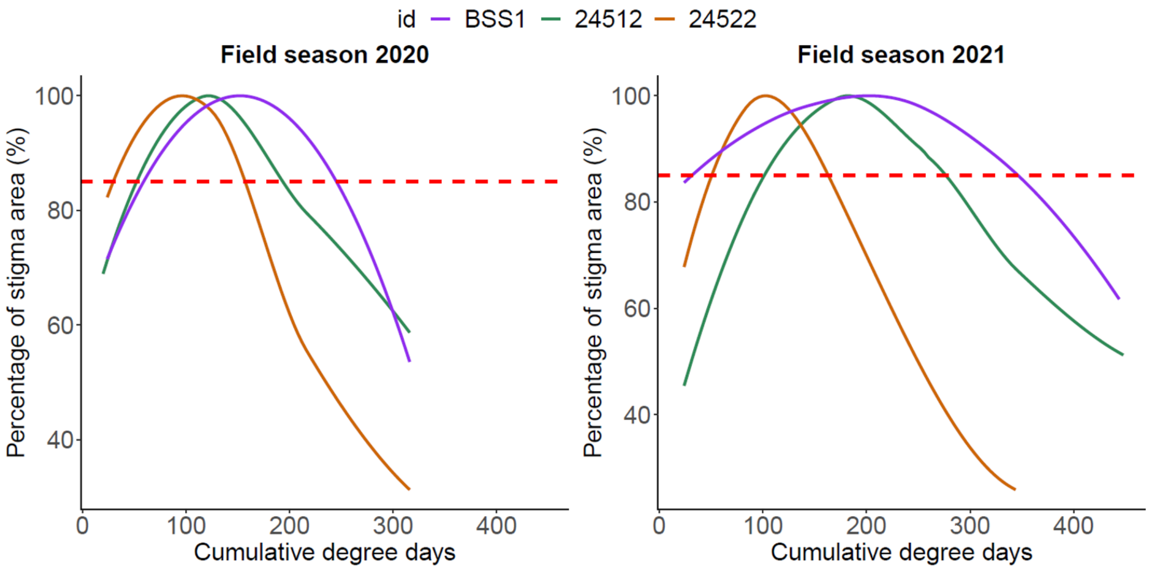


**Figure S6. Developmental stigma patterns expressed in percentage from maximum observed stigma area.** Red dashed line indicates 85% benchmark for the selection of the boundaries of the peak phase with the growth and deterioration phases.

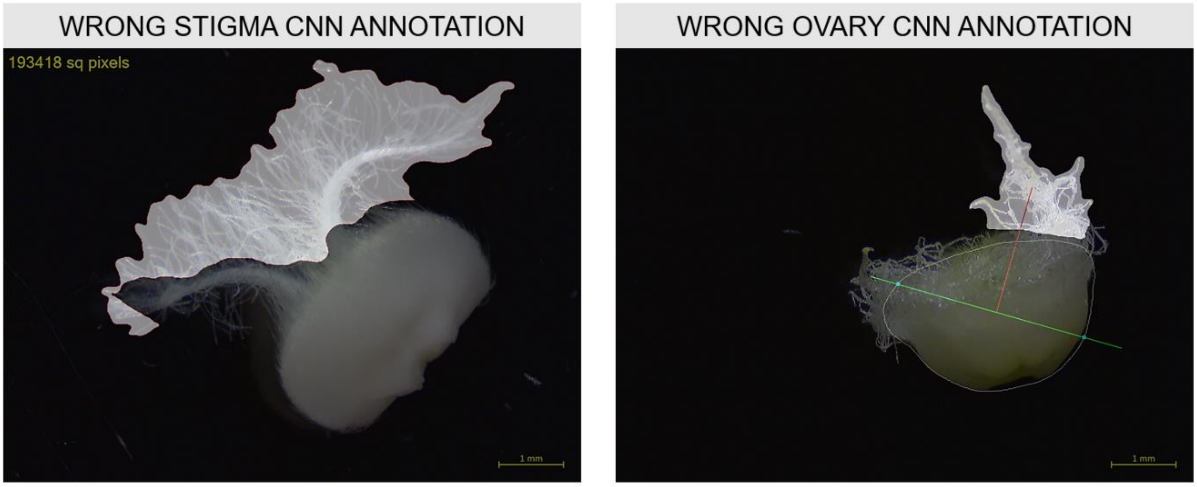


**Figure S7. Erroneous stigma area and ovary diameter CNN annotations.** Examples of bright field images of carpels wrongly annotated by the stigma (left) and ovary (right) CNNs.
